# Slow RNAPII elongation enhances naïve-pluripotency rewiring while preserving replication fork speed

**DOI:** 10.1101/2025.05.12.653265

**Authors:** Sara Martín-Vírgala, Joana Segura, Alicia Gallego, Javier Isoler-Alcaraz, Lothar Schermelleh, María Gómez

## Abstract

DNA replication and transcription must be intricately coordinated, as both machineries navigate the same chromatin landscape to ensure genome stability and proper cell function. Here, we uncover that a global imbalance between their elongation rates— specifically, slowed transcriptional elongation alongside rapid replication fork progression—does not elicit replicative stress. Instead, this uncoupling accelerates the acquisition of naïve pluripotency during *in vitro* de-differentiation, revealing an unexpected link between transcription kinetics and cell plasticity. Mechanistically, we show that the transition to naïve pluripotency is accompanied by a distinctive alternative splicing program indicative of reduced RNAPII elongation, both *in vitro* and *in vivo*. These findings redefine the functional relationship between replication and transcription dynamics and uncover transcriptional velocity as a tunable layer of control over cellular identity transitions.

**Highlights:**

1. Replication and transcription elongation rates can be uncoupled genome-wide.
2. Slow transcription elongation accelerates the acquisition of naïve pluripotency during *in vitro* de-differentiation.
3. High replication fork speed is maintained in slow-transcribing cells during cell state transitions.
4. Alternative splicing is distinctly regulated at the naïve and primed pluripotency states, both *in vitro* and *in vivo*.

## INTRODUCTION

DNA replication and transcription are two essential processes that utilize the genome as a template. Accumulated evidence over the last decades showed that conflicts between them threaten genome stability (Goehring et al., 2023; Lalonde et al., 2021; Gómez-González and Aguilera, 2019; Hamperl and Cimprich, 2016). To minimize the potentially detrimental collisions between translocating replication and transcription complexes, cells have evolved a myriad strategies. This includes co-orienting the directionality of both processes, most evident in bacteria (Browning and Merrikh, 2024), initiating replication close to transcription start sites (TSS) of genes (Chen et al., 2019; Lombraña et al., 2013; Cayrou et al., 2011; Sequeira-Mendes et al., 2009), using on-going transcription to push replicative helicases away from the gene bodies (Liu et al., 2021; Lombraña et al., 2016; Gros et al., 2015; Powell et al., 2015), or even delaying the duplication of the TSS of active genes till very late in the cell cycle (Wang et al., 2021). Beyond this, recent research unveiled the tight relationship between both processes when they co-occur in the genome, as shown by the fast re-binding of active RNA polymerase II (RNAPII) complexes onto replicated DNA minutes after the passage of the replication fork (Bruno et al., 2024; Fenstermaker et al., 2023). This coupling might facilitate a rapid re-establishment of the transcriptional program after DNA replication, thus contributing to maintain cellular identity (Fenstermaker et al., 2024, 2023).

A yet unexplored angle of the crosstalk between both processes pertains to the speed at which they occur in the genome. In bacteria, the replication rate is around 12 times faster than the transcription rate (600 nt/sec vs 50 nt/sec, respectively) (Helmrich et al., 2013). Thus, the divergent replication forks emanating from the single bacteria replication origin would unavoidably collide with transcribing polymerases. The fact that the large majority of highly expressed and essential genes are co-directionally oriented relative to the replisome across bacterial species limits the potentially more harmful head-on encounters (Rocha and Danchin, 2003; Zheng et al., 2015). In contrast, in the large genomes of mammalian cells, where DNA replication initiates from thousands of replication origins, the speed of elongating RNAPII and replication forks fells within a similar range (17-33 nt/sec *vs* 17-72 nt/sec, respectively) (Helmrich et al., 2013). Similar translocation rates, in principle, might help minimize the encounters between both machineries. Indeed, replication-transcription conflicts do occur at long genes that need more than one cell cycle to be transcribed, contributing to the instability of common fragile sites within those genes (Blin et al., 2019; Helmrich et al., 2011).

Here, we set out to investigate the relevance of balanced replication-transcription rates for genome function. For that, we employed mouse embryonic stem cells (mESC) carrying a single residue substitution in RPB1, the largest subunit of RNAPII (Maslon et al., 2019). The R749H mutation in RPB1 impairs early mouse development, but mESCs can be maintained in culture and display a slow transcription elongation phenotype (from an average of 2,450 nt/min in WT cells to 1,780 nt/min in RNAPII-mut cells, respectively; Maslon et al., 2019). By combining time-resolved analyses of transcriptome rewiring and replication dynamics upon cell de-differentiation, we show that slow RNAPII transcription promotes naïve-pluripotency while maintaining fast replication fork speed. Moreover, we found that unbalanced transcription and replication rates do not generate replicative stress, suggesting that modulating RNAPII velocity could provide new strategies to enhance cell plasticity and reprogramming.

## RESULTS

### Slow transcription elongation rate leads to fast replication fork rate and restrains replicative stress

To investigate the impact of slow transcriptional rates on DNA replication, we employed mESCs knocked-in for a slow RNAPII mutant (R749H) generated at the Cáceres lab (Maslon et al., 2019). Replication elongation rates were addressed by stretched DNA fiber assays (Figure 1A), which showed that RNAPII-mut cells display significantly faster replication forks than their WT counterparts (Figure 1B). This increase in fork speed, however, does not lead to a compensatory effect on the number of active replication origins (Figure 1C), nor affects the symmetrical progression of the forks (Figure 1D). Measuring global DNA synthesis rates by EdU incorporation either by immunofluorescence or flow cytometry confirmed the enhanced replication of RNAPII-mut cells (Figure 1E and Figure S1A). Importantly, increased replication rates did not alter the overall duration of the cell cycle nor the proportion of cells in the different sub-stages of the S-phase (Figure S1A-C). Consistent with unperturbed DNA replication progression, RNAPII-mut cells do not show signs of DNA damage or replication stress (Figure S1D). On the contrary, slower transcription and faster replication results in slightly reduced levels of γH2AX signaling (Figure S1D-E), more evident at late S-phase (Figure 1F). This result was not due to RNAPII-mut cells having an impaired response to DNA damage, since γH2AX signalling was readily increased when cells underwent replicative stress upon aphidicolin exposure or DNA damage by γ-irradiation (Figure S1D and S1F). These observations suggest that reducing RNAPII speed allows faster moving forks and limits replicative stress.

**Figure 1.**
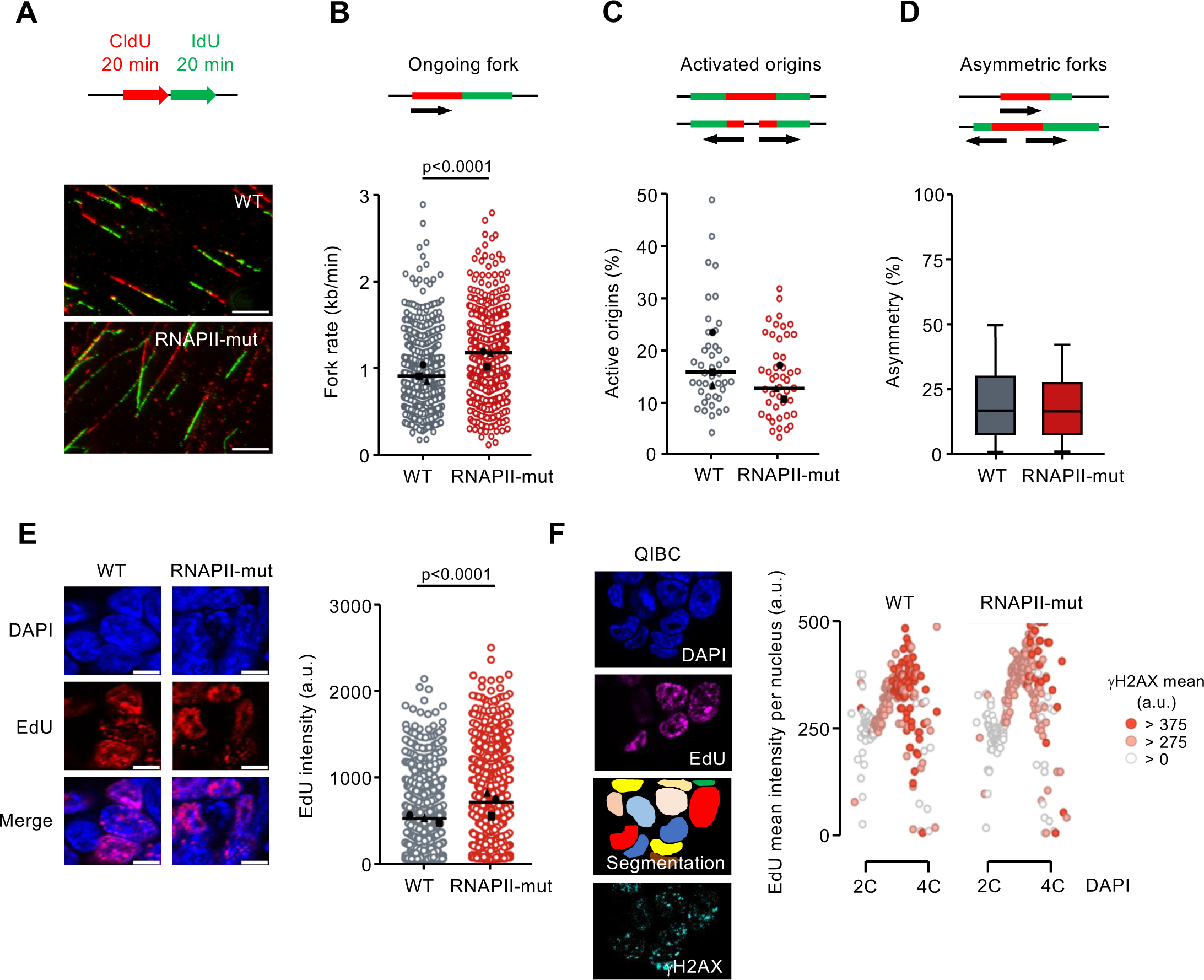
Slow transcription elongation rate leads to fast replication without generating replicative stress. (A) Experimental design and representative images of stretched DNA fibers from WT and RNAPII-mut cells. Scale bar 10 μm. (B-D) Dot plots displaying the fork rate (B), first-labelled origins (FLO, C) and fork asymmetry (D) from WT and RNAPII-mut cells. Median values for each replicate are indicated with a black symbol (n=3) and the global median with a horizontal line. n>669 tracks for FR and n>44 images for FLO calculations, respectively. Mann-Whitney test significant p-values are shown on top, as well as a scheme of the replication parameters measured in each case. (E) Representative confocal microscopy images showing EdU Click-It signal in WT and RNAPII-mut cells. Cells were pulse-labelled with 10 μM EdU for 20 min before fixation. Scale bar 5 μm. Data are from 3 independent experiments. Mean values are indicated as in B-C. WT n=1405, RNAPII-mut n= 1169. Mann-Whitney test significant p-values are shown on top. (F) Quantitative Image Based Cytometry (QIBC) scatterplots of DNA content (DAPI) versus EdU signal, showing γH2AX nuclear intensity in colour scale at the different cell cycle phases. A single representative experiment is shown, with n=18.000 analysed nuclei per condition. Nuclei displayed on the graphs have been grouped according to DAPI intensity intervals of 5,000 units (binning).

Faster fork rates with a similar number of active origins should shorten S-phase length. To determine if this was the case, cells were synchronized at the G1/S border by a double thymidine block and DNA synthesis along S-phase was monitored by flow cytometry (Figure S2A). RNAPII-mut cells enter faster in early-S and progress faster to late-S (Figure S2A-B, see time points 1.5h and 4.5h after thymidine-release), altogether resulting in a slightly shorter S-phase. To address whether faster fork rates were constant along S-phase, we performed DNA fiber analyses at 1h, 3.5h and 6h after thymidine-release, representing early, mid and late S-phase time points (Figure S2C). We found that fork rate accelerates along S-phase, in agreement with recent work employing scEdU sequencing in HeLa cells (van den Berg et al., 2024). Despite some differences at the late-S time point, fork dynamics along S-phase were overall comparable between WT and RNAPII-mut cells.

### Replication-transcription encounters remain similar despite differences in transcription and elongation rates

We wondered whether the fast replicative phenotype of RNAPII-mut cells arose from reduced encounters between the replication and transcription machineries. Since the equivalent R749H mutation in *Drosophila* shows a diminished ability to read through intrinsic elongation blocks (Chen et al., 1996), which might alter its processivity on chromatin, we first quantified the number of active transcriptional complexes in S-phase. 3D-SIM superresolution microscopy (Figure S2D), as well as chromatin fractionation analyses (Figure S2E), showed that both WT and RNAPII-mut cells have similar number of elongating transcriptional complexes (RNAPII phosphorylated on Ser2 of its C-terminal domain, RNAPII S2P) in early and mid S phase. We then measured the co-localization of RNAPII S2P with sites of on-going replication. To maximize the resolution of our analyses cells were pulse-labeled with EdU for 5 min (labeling nascent DNA fragments of around 5 kb in euchromatic replication) (Figure 2A). We found that the percentage of overlapping EdU/RNAPII S2P foci per nucleus decreased from early to mid S-phase (Figure 2B), but was comparable between cell types at both sub-S phases. Notably, the average overlap between elongating replication and transcription sites per nucleus match previous estimations by proximity ligation assays (PLA) in human cell lines (Fenstermaker et al., 2023). We speculate that the reduced cell-to-cell variation in the co-localization frequency of replication and transcription sites in early-S RNAPII-mut cells could result from their more homogeneous and slower transcription elongation rates (Maslon et al., 2019).

**Figure 2.**
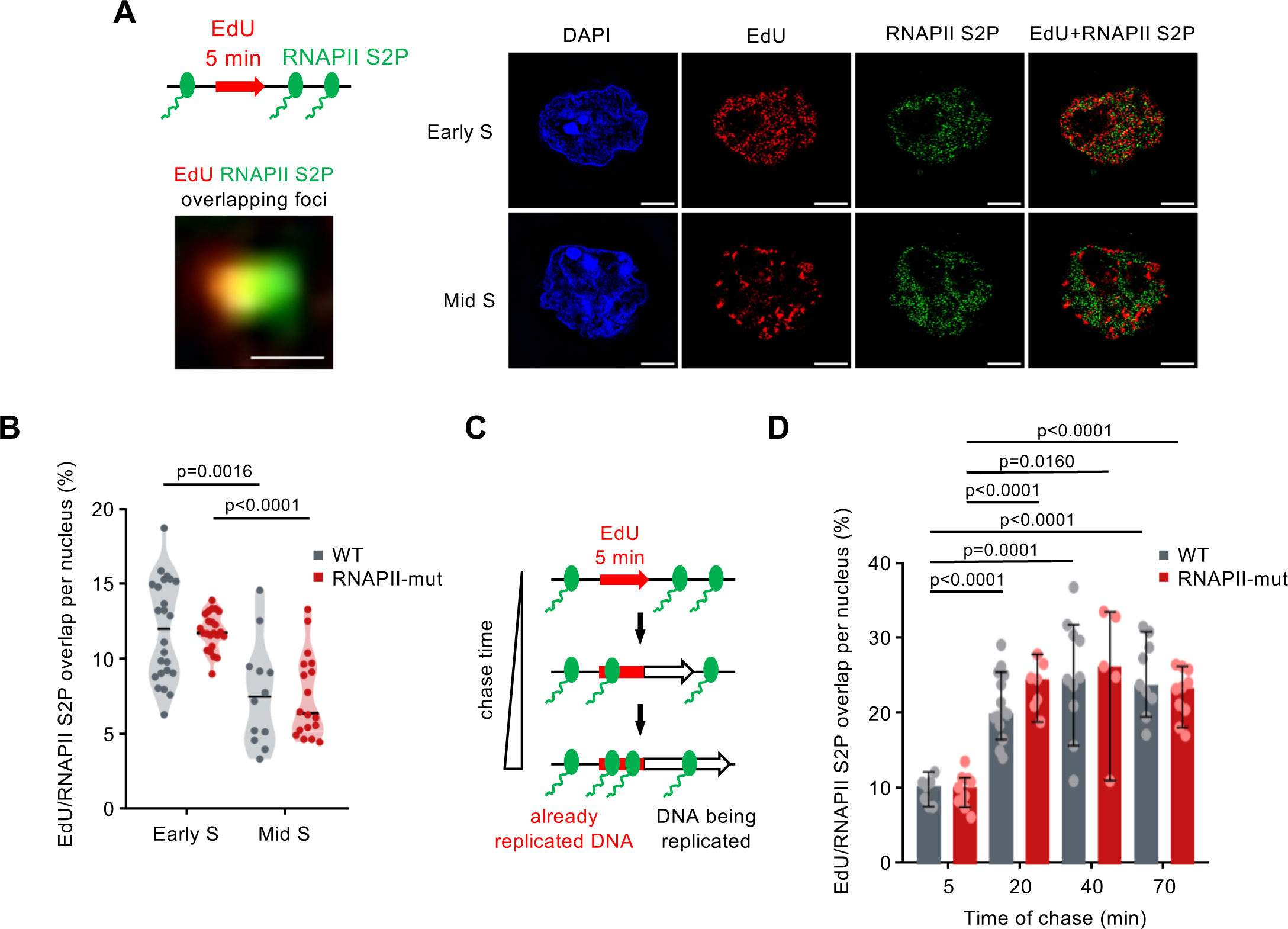
The number of encounters between active replication and transcription complexes is similar in WT and RNAPII-mut cells. (A) Experimental scheme and representative 3D-Structured-Illumination Microscopy (3D-SIM) images of a single xy plane of WT cells in early and mid S-phase according to their EdU patterns, together with RNAPII S2P foci location. Scale bar 4 μm. On the bottom left, representative zoomed image of the overlapping RNAPII S2P/EdU foci quantified in B and D. Scale bar 0.2 μm. (B) Violin plots showing the percentage of RNAPII S2P foci overlapping with EdU per nucleus, normalized by the total number of RNAPII S2P foci in each nucleus. Median values are indicated by a horizontal line. Data are pooled from 2 independent experiments. Mann-Whitney test significant p-values are shown on top. Number of nuclei: WT early n=24, RNAPII-mut early n=23, WT mid n=12, RNAPII-mut mid n=18. (C) Schematics of the pulse-chase experimental design to assess RNAPII S2P reloading on post-replicative chromatin in early S phase. (D) Graph showing the percentage of RNAPII S2P foci overlapping with EdU per nucleus, normalized by the total number of RNAPII S2P foci in each nucleus at each chase time point. Medians and 95% CI (Confidence Interval) are shown. Data shown are from a single representative experiment. Number of nuclei: WT chase 5 min=7, RNAPII-mut chase 5 min=9, WT chase 20 min=14, RNAPII-mut chase 20 min=7, WT chase 40 min=10, RNAPII-mut chase 40 min=5, WT chase 70 min=9, RNAPII-mut chase 70 min=10. Welch’s test was used to assess statistical significance between groups. P-values <0.05 are shown on top.

To determine if the R749H mutation leads to an altered recruitment of RNAPII to nascent DNA, cells were labelled with EdU for 5 min followed by increasing chase times to assess RNAPII S2P occupancy following DNA replication (Figure 2C). We observed significant increases in the percentage of overlapping EdU/RNAPII S2P foci from 20 min chase time, reaching their maximum at 40 minutes after replication (Figure 2D). These results confirm recent findings showing that the elongating form of RNAPII associates with DNA within a few minutes after the passage of the replication fork (Bruno et al., 2024; Fenstermaker et al., 2023). As before, no significant differences were found between WT and RNAPII-mut cells, indicating that the dynamics of RNAPII recruitment onto replicated DNA were similar in both cell types. Thus, contrary to our expectations, faster replication fork rates in RNAPII-mut cells were not due to reduced encounters between replication and transcription complexes on chromatin.

### RNAPII-mut cells reach a fully naïve-pluripotency state without reducing replication fork rates

mESCs in culture fluctuate between multiple interconvertible states and this metastability and plasticity is thought to be at the basis of their pluripotent identity (Kumar et al., 2014; Cahan and Daley, 2013; Loh and Lim, 2011; Martinez Arias and Brickman, 2011). To understand the functional implications of slow transcription elongation on cell plasticity we challenged WT and RNAPII-mut mESCs to undergo *in vitro* de-differentiation by the addition of MEK and GSK3 inhibitors (two-inhibitor cocktail or 2i) to the culture medium. This well-established 7-days protocol insulates cells from differentiation stimuli shifting the population towards a more undifferentiated state known as naïve-pluripotency (Hackett and Surani, 2014; Nichols and Smith, 2009). For simplicity, we refer to the cells at day 0 as primed (cells grown in serum+Lif; primed-pluripotency state) and to the cells at day 7 as naïve (cells grown in serum+Lif+2i; naïve-pluripotency state) (Figure 3A). After 7 days of 2i exposure, both WT and RNAPII-mut cell colonies acquire a rounder morphology and increase the expression of the pluripotency factor NANOG, while reducing the expression of the DNA methyltransferase DNMT3b (Figure 3B-C), as reported (Ubieto-Capella et al., 2024; Martinez-Val et al., 2021; Lynch et al., 2020). To evaluate whether RNAPII-mut cells fully attain the naïve-pluripotency identity we performed RNA-seq to profile gene expression along the de-differentiation regime (days 0, 1, 2 and 7 of 2i addition). Differential expression analyses between RNAPII-mut and WT cell types at primed-pluripotency state (day 0) identified 306 up and 393 down differential expressed genes (DEGs), respectively (Figure S3A). Notably, downregulated genes exhibit greater fold-change values than upregulated ones, which might be related to the slow elongation rate phenotype of these cells. Consistent with this interpretation, downregulated genes are significantly longer in size than the average (Figure S3B), in line with the reported downregulation of long synaptic genes in RNAPII-mut neuronal progenitors (Maslon et al., 2019). This was expected since longer genes tend to be more prone to RNAPII failure during elongation. In addition, downregulated genes are little expressed even in WT conditions (Figure S3C), suggestive of fewer numbers of elongating RNAPII molecules traversing these genes. While downregulated DEGs account for genes related to cell migration and extracellular matrix organization, the upregulated ones are enriched in terms related to translation (Figure S3D). Despite these differences in gene expression between WT and RNAPII-mut cells at the primed-pluripotency state, the magnitude of expression changes along the de-differentiation process was strongly correlated between cell types (Figure 3D). In agreement, both cell types displayed parallel patterns of expression of key pluripotency genes (Figure S3E). Moreover, Principal Component Analysis (PCA) indicates that, although through different trajectories, WT and RNAPII-mut cells achieve a similar naïve-pluripotency stage at day 7 (Figure 3E). These results imply that altering transcription and replication elongation rates do not hamper mESCs plasticity.

**Figure 3.**
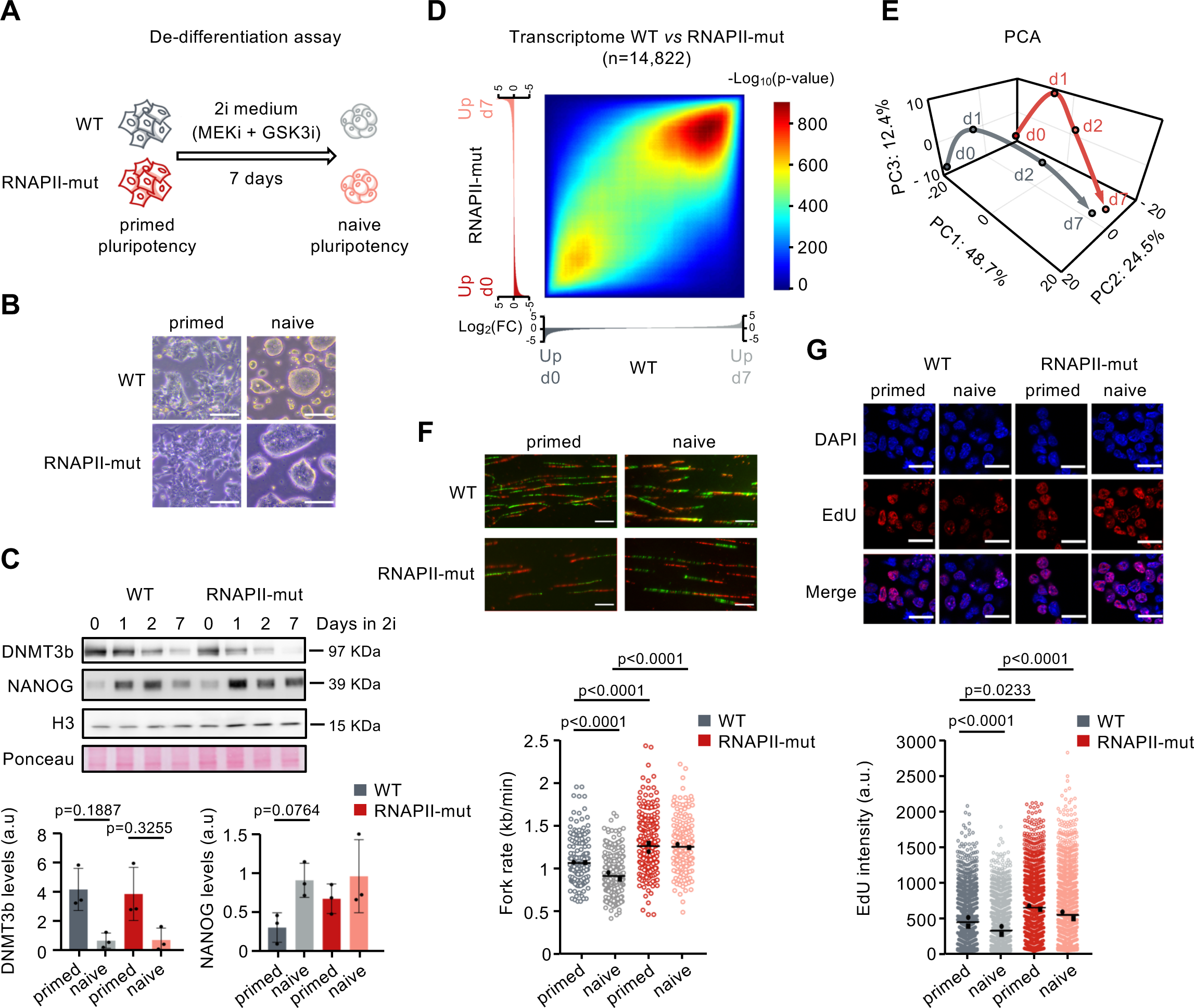
RNAPII-mut cells reach a fully naïve-pluripotency state without reducing replication fork rates. (A) Scheme of the primed-to-naïve pluripotency de-differentiation assay. (B) Brightfield representative images showing the morphological changes after 7 days of 2i-exposure in WT and RNAPII-mut cells. Scale bar, 100 µm. (C) Top, representative immunoblot of whole cell extracts showing NANOG and DNMT3b levels during the indicated days of the de-differentiation regime. Total histone H3 levels and Ponceau staining were used as loading controls. The size of the detected proteins is indicated on the right. Bottom, histograms showing NANOG and DNMT3b levels from three independent experiments (black dots) for WT and RNAPII-mut cells in primed (day 0) and naïve (day 7) states. Statistical significances were assessed by Krustkal-Wallis test. P-values are indicated (D) Rank-Rank Hypergeometric Overlap (RRHO) plot showing the significance of the overlaps between ranked sets of differentially expressed genes (day 7 versus day 0) in WT and RNAPII-mut cells (2 biological replicates; n=14,822 genes). First, genes were arranged by their log_2_(fold-change), then assessed for overlap by RRHO sliding window of 100 genes. Color intensity represents -log_10_(p-value) after Benjamini-Yekutieli correction of hypergeometric overlap. (E) Principal Component Analysis (PCA) based on gene expression in WT (grey) and RNAPII-mut (red) cells grown for 0, 1, 2, and 7 days in 2i medium. Dots represent the merge of two replicates in each condition. First three principal components are shown. (F) Representative images of stretched DNA fibers and dot plots showing fork rate (FR) measurements in primed-pluripotency and naïve-pluripotency WT and RNAPII-mut cells. Scale bar, 10 μm. Symbols indicating median values of the individual replicates (n=2) and total medians are as in Figure 1B. n>152 structures/condition. Mann-Whitney test significant p-values are shown on top. (G) Representative confocal images and quantification of EdU Click-It signal intensity per nucleus in primed and naive WT and RNAPII-mut cells. Nuclear DNA is stained with DAPI. Scale bar, 15 μm. Data are for 2 independent experiments. Mean values are indicated as in Figure 1F. Number of nuclei: WT primed=1893, WT naïve=2272, RNAPII-mut primed=1263, RNAPII-mut naïve= 1400. Statistical analysis was conducted with Krustkal-Wallis test. P-values are indicated.

We next addressed replication fork rates in naïve cells. Previous studies have shown that the 2i de-differentiation course induces replication fork slowdown in mESCs (Ubieto-Capella et al., 2024), resembling the situation in totipotent-like cells (Nakatani et al., 2022) and in preimplantation embryos, where the speed of the forks is very slow at the initial stages and increases along development (Takahashi et al., 2024; Nakatani et al., 2022). Surprisingly, while WT cells recapitulate fork slowdown during the primed-to-naïve transition, RNAPII-mut cells remained fast replicating (Figure 3F). A similar result was obtained by measuring global DNA synthesis rates by EdU incorporation (Figure 3G). To get a mechanistic insight for this phenotype, we checked the expression levels of the ribonucleotide reductase (RNR) subunits RRM1 and RRM2, the enzymes mediating deoxynucleotide triphosphate (dNTP) production required to sustain DNA synthesis (Fairman et al., 2011) (Figure S3F). Notably, both *Rrm1* and *Rrm2* genes become downregulated along the de-differentiation course in WT cells but not in RNAPII-mut ones, mirroring the dynamics of DNA replication in each cell type. This indicates that the expression of RNR subunits is regulated during pluripotency transitions, but RNAPII slow elongation overrides the need for this regulation. These findings demonstrate that replication and transcription rates can be globally uncoupled.

### Changes in transcript isoforms suggest slower transcription elongation speeds at earlier cell and embryonic pluripotency stages

Optimal transcription elongation rates are required to ensure proper alternative splicing (AS), a key process that enhances RNA diversity contributing to protein isoform variation (Maslon et al., 2019; Saldi et al., 2016; Jimeno-González et al., 2015; Braunschweig et al., 2014; Dujardin et al., 2014; Fong et al., 2014; Ip et al., 2011; de la Mata et al., 2003). Changes in AS, in turn, can influence the outcome and fate of transcripts, constituting an additional layer of regulation of cellular states (Baralle and Giudice, 2017). We therefore set out to determine the AS profiles in naïve and primed pluripotency stages and use them to infer RNAPII speed changes at this early cell transition window.

First, we compared the total number of AS events in primed WT and RNAPII-mut cells using *vast-tools*, which assigned a PSI value (Percent Spliced-In) to each specific event (Figure 4A and S4A). As expected, we found a transcript-wide increase in the number of AS events in slow-elongating RNAPII cells, in particular higher numbers of spliced introns (Figure 4B). This pattern was similarly observed at the original dataset characterizing the slow transcriptional speed of the R749H mutation, although the cells were grown in a different media there (serum + 2i, no Lif; Maslon et al., 2019) (Figure 4C). We then computed the number of differential splicing events (ΔPSI) between both cell types and detected 427 differentially regulated introns. Of those, RNAPII-mut cells displayed higher levels of intron retention compared to WT cells, a feature that was also present in the other dataset (Figure 4D and S4B). Therefore, we defined the AS signature associated with slow transcription elongation in this cellular stage as a gain in intron retention events.

**Figure 4:**
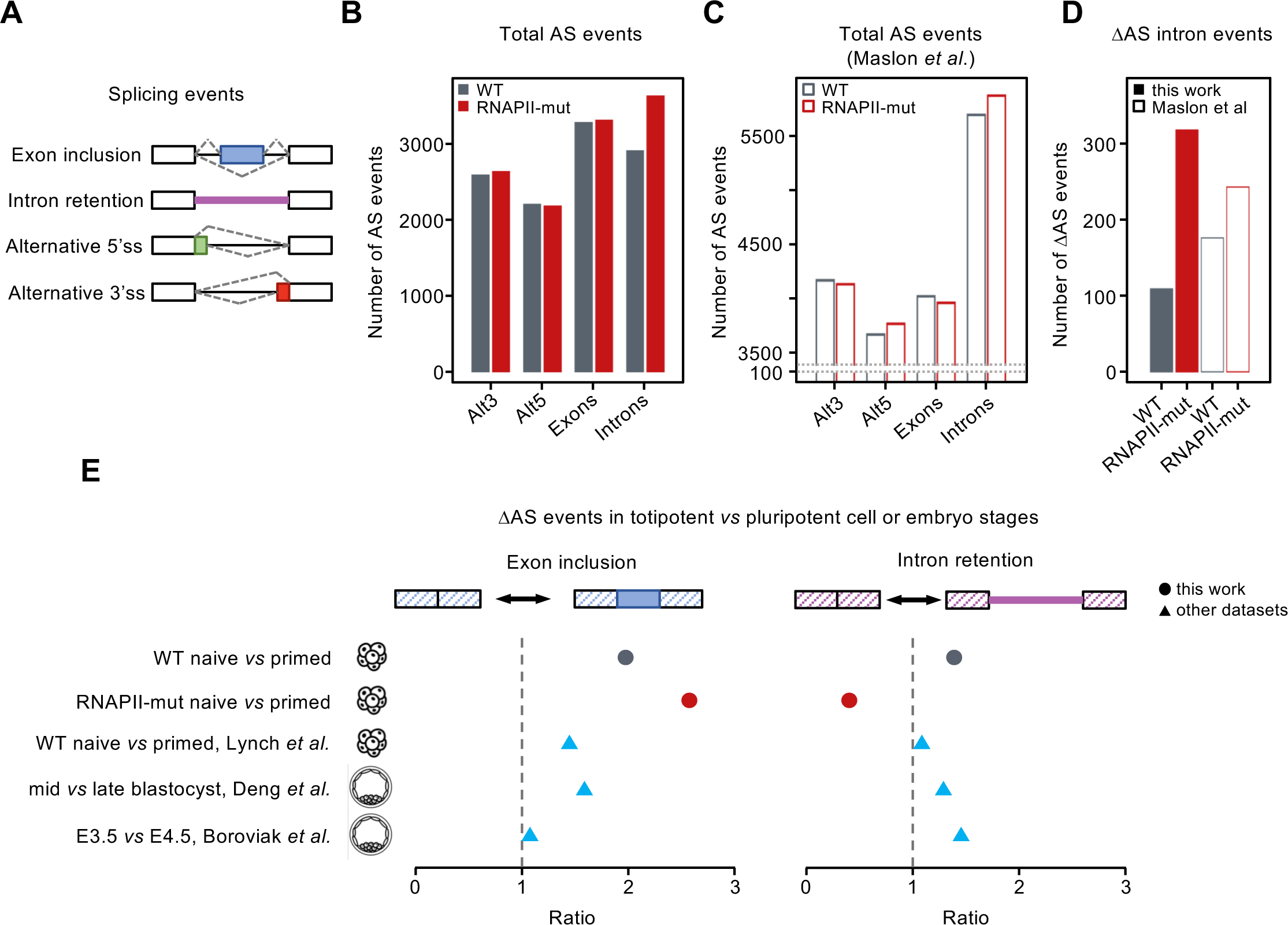
Changes in transcript structure at naïve and primed-pluripotency cells and embryonic stages. (A) Schematic representation of the alternative spliced (AS) events (PSI values between 10-90%) analysed. (B-C) Total number of AS events detected in merged replicates of WT and RNAPII-mut cells at primed state (B), and in Maslon *et al* (2019) dataset (C), where cells were grown in serum + 2i. (D) Number of differentially retained introns in RNAPII-mut *vs* WT in our dataset (full bars) and Maslon *et al* (2019) dataset (empty bars). (E) Differences in exon inclusion (left) and intron retention (right) levels between totipotent and pluripotent cells and embryonic stages in the indicated datasets. Data from Lynch *et al* (2020) correspond to WT mESCs grown in similar conditions to our work (serum + Lif -/+ 2i). Data from Deng *et al* (2014) and Boroviak *et al* (2015) correspond to *in vivo* samples from the indicated mouse embryo developmental states. Values on the right side of the dotted line indicate higher exon inclusion or intron retention in the more totipotent stages.

We next quantified the ΔPSI between naive and primed pluripotency states in WT cells. This analysis identified a tendency towards increasing exon inclusion and intron retention events in naïve samples (Figure 4E and S4C-D), indicative of a distinct regulation of AS in different pluripotency stages. This trend was confirmed in datasets from published mESC de-differentiation course (Lynch et al., 2020) and, most importantly, in datasets from mouse embryos spanning comparable developmental period (E3.5 *vs* E4.5 or mid-*vs* late-blastocist, respectively) (Boroviak et al., 2015; Deng et al., 2014). Strikingly, while RNAPII-mut cells similarly display increased ratios of exon inclusion in naïve *vs* primed pluripotency stages (Figure 4E, left plot), they show the opposite tendency of intron retention (Figure 4E, right plot). Since RNAPII-mut cells already displayed intron retention gains in the primed-pluripotency state, this result suggests that RNAPII elongation rate is slower at naive-pluripotency than at primed-pluripotency in WT cells and embryos.

### Slow transcription elongation accelerates the transcriptional switch towards naïve-pluripotency

The above results postulate that slow transcription elongation rate could be a trait of early pluripotency stages. This hypothesis aligns with the observations that (i) RNAPII-mut cells expressed higher levels of the pluripotency marker NANOG at primed state (Figure 3C), and that (ii) they appear more advanced towards the stabilization of the naïve-pluripotency state upon de-differentiation (Figure 3E). To unveil the molecular basis underlying this effect we generated gene regulatory modules (UMAPs) based on normalized expression values (Z-scores) of the differentially expressed genes in at least one comparison along the WT cells de-differentiation scheme (n=1878) (Figure 5A). Genes clustered in 4 groups reflecting their expression trajectories across the time-course: gradual or rapid decreases (clusters 1 and 2, respectively) and rapid or gradual increases (clusters 3 and 4, respectively) (Figure 5B and S5A). According to previous characterizations of the 2i-induced primed-to-naïve transition, GO-terms analyses of these clusters revealed the silencing of genes related to multicellular organism development, cell differentiation and adhesion, and the upregulation of genes involved in lipid metabolism and catabolic processes (Figure S5B) (Martinez-Val et al., 2021; Ubieto-Capella et al., 2024).

**Figure 5.**
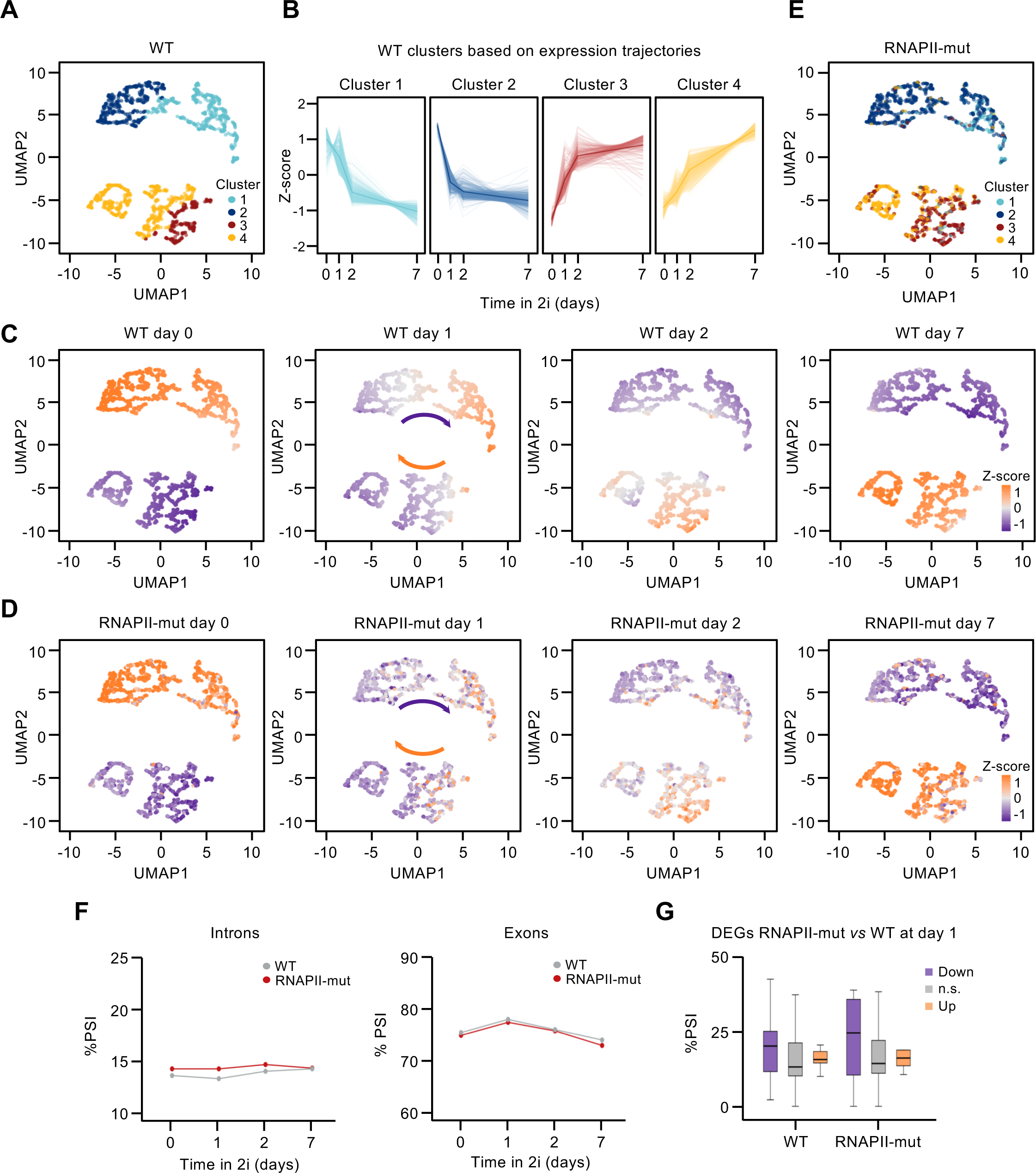
Slow transcription elongation accelerates the transcriptional switch towards naïve-pluripotency. (A) UMAP analysis representing the similarity of coding genes in terms of their expression levels (Z-scores) in WT cells, considering the set of genes that are differentially expressed (DEGs) in at least one naïve versus pluripotent comparison (d1-d0, d2-d0 or d7-d0) (n=1878 genes). Colors represent gene clusters identified by hierarchical clustering of their expression trajectories across WT dedifferentiation (Figure 5SA). (B) Trajectories of normalized Z-score expression levels across WT cells dedifferentiation (from d0 to d7) of DEGs shown in A. Cluster 1 (gradual decrease) n=517; cluster 2 (rapid decrease) n=460; cluster 3 (rapid increase) n=267; cluster 4 (gradual increase) n=634. (C-D) UMAP genes shown in A colored based on normalized Z-score expression levels at the four time points d0, d1, d2, d7 in WT cells (C) or RNAPII-mut cells (D). (E) UMAP genes shown in A, colored based on the hierarchical clustering of their expression trajectories across RNAPII-mut cells de-differentiation. (F) Median PSI values of the indicated AS events in UMAP genes shown in A along the primed-to-naïve transition. (G) PSI distributions of AS events in UMAP genes shown in A sorted by their differential expression in the RNAPII-mut *vs* WT comparison at day 1 of dedifferentiation, as indicated.

As anticipated by the PCA analyses, projecting gene expression levels along the de-differentiation course in WT and RNAPII-mut cells revealed that the main differences between cell types were detected at day 1, where more genes transition to increased or decreased expression in the latter (Figure 5C-D, panels for day 1). Projection of the gene clusters defined in RNAPII-mut cells on the WT-UMAPs further show that this observation accounts for changes in gene clusters (i e : cluster 2 genes - rapid decrease-occupy the space of cluster 1 genes -gradual decrease- and the opposite; cluster 3 genes -rapid increase-occupy the space of cluster 4 ones -gradual increase-) (Figure 5E). Nevertheless, and in agreement with the similar naïve-pluripotency stage reached by both cell types, the 4 gene clusters defined in RNAPII-mut cells comprise similar functional categories to the WT ones (Figure S5C). These findings indicate an accelerated reprogramming of gene expression in RNAPII-mut cells.

Although expression changes and AS changes did not correlate (Figure S4E and Maslon et al., 2019), we wondered whether the AS patterns of these UMAP DEGs vary along the de-differentiation program. Interestingly, DEGs in WT cells showed a tendency towards increasing intron retention over time, reaching a similar level to RNAPII-mut cells at the naïve stage (Figure 5F, left panel). In contrast, exon inclusion levels were parallel in both cell types at all time points (Figure 5F, right panel). This behavior indicates that AS patterns are dynamic during primed-to-naïve de-differentiation in WT cells and suggests that, at naïve stages, WT cells displayed an AS signature associated with slow transcription elongation, as evidenced by their increased intron retention levels. Supporting this, the genes that respond faster to the de-differentiation cues in RNAPII-mut cells (the temporarily-differential genes at day 1), already showed a tendency to gain intron retention events in WT cells, although not as marked as in their mutant counterparts (Figure 5G). Altogether, we conclude that slow RNAPII elongation favors a faster transcriptional rewiring towards naïve-pluripotency.

## DISCUSSION

Here we analysed the consequences of slow transcription elongation for DNA replication and stem cell plasticity. First, we found that slow transcription leads to fast replication without generating conflicts between both processes. Moreover, this speed unbalance does not prevent stem cells from transitioning efficiently to a naïve-pluripotency state. On the contrary, slow transcribing cells are predisposed to transcriptome rewiring towards naïve-pluripotency and they do so while maintaining fast replication rates. Second, we provide evidence arguing that the naïve-pluripotency state in mESCs and embryos associates with slower RNAPII transcription rates. These observations demonstrate that replication and transcription rates can be uncoupled in stem cells and suggest an unexpected link between RNAPII elongation speed and cell plasticity, providing insights for future interventions manipulating transcription velocity towards enhancing reprogramming.

How can slow transcription elongation lead to fast replication elongation? The simplest explanation is that it increases the availability of ribonucleotides, precursors of the building blocks for replication. Indeed, nucleotide depletion reduces replication fork velocity and *vice versa* nucleoside supplementation accelerates fork speed (Somyajit et al., 2017; Poli et al., 2012). In agreement with this, the expression levels of *Rrm1* and *Rrm2* were slightly lower in RNAPII-mut that in WT cells at the primed state, suggestive of higher ribonucleotide pool in the former. Since accelerating fork speed rescues replicative stress in mouse and human embryonic stem cells (Halliwell et al., 2020; Almeida et al., 2018), another possibility is that the fast replication of RNAPII-mut cells arises as a compensatory response to limit transcription-mediated replicative stress. The unexpected lack of increased conflicts between replication and transcription in slow-elongating RNAPII cells aligns with this interpretation.

The molecular basis of slow fork rates in developing mouse embryos as recapitulated *in vitro* in naive-pluripotency stages is not yet fully understood. Slower forks might contribute to higher replication fidelity at early stages to cope with the structural and epigenetic reprogramming occurring in mammalian embryogenesis (Pal et al., 2025; Ke et al., 2017) or to facilitate key transcriptional changes accompanying embryonic development (Nakatani et al., 2022). However fork acceleration by nucleoside supplementation in early embryogenesis is generally error-free (Takahashi et al., 2024), arguing against fidelity being the constraint for regulating replication fork velocity. Similarly, whether slow RNAPII elongation contributes to transcriptional fidelity at early developmental stages is unknown. In a recent report, Debés and collaborators (2023) showed that overall RNAPII speed increases with age in different organisms and tissues, and that these changes in elongation speed were accompanied by changes in splicing. Most interestingly, they demonstrated that worms and flies carrying a mutated RNAPII orthologous to the R749H studied here have an increased lifespan, suggesting that slow RNAPII elongation supports transcriptional accuracy (Debès et al., 2023). We propose that RNAPII elongation speed is regulated at the transition through pluripotency, and might underlie the acquisition of the transcriptional identity occurring at this developmental period, when the lineage decision specifying epiblast and primitive endoderm cell fate choice occurs (Bassalert et al., 2018; Boroviak and Nichols, 2014; Ohnishi et al., 2014). Further *in vivo* studies are required to test this hypothesis.

Slow RNAPII elongation is also linked to the enhanced co-transcriptional deposition of N6-methyladenosine (m6A) (Gallego et al., 2022; Slobodin et al., 2017), the most abundant internal mRNA modification in eukaryotes that is involved in several key RNA processes, including splicing, stability and translation (Louloupi et al., 2018; Barbieri et al., 2017; Ke et al., 2017). Interestingly, m6A reduces RNA stability in mESCs (Liu et al., 2020), which might be related to the observed accelerated silencing of key genes during the de-differentiation process of RNAPII-mut cells. In addition, the increased intron retention shown by these transcripts might lead to their degradation via nonsense-mediated decay (Braunschweig et al., 2014). Understanding how RNAPII elongation rate, m6A regulation and alternative splicing dynamics converge to fine-tune the regulation of naïve and primed pluripotency is an exciting avenue for future research.

### SUPPLEMENTARY FIGURE LEGENDS

**Figure S1 (related to Figure 1). Cell cycle characterization and DNA damage response in WT and RNAPII-mut cells.** (A) Left, representative flow cytometry dot plots showing the different cell sub-populations according to their cell cycle stage. Cells were labelled with 10 μM EdU for 30 min. Right, quantification of the percentage of cells in each cell cycle phase. Mean and SD are shown. Data are from 3 independent experiments. Statistical significance was analysed by a Student t-test comparing WT *versus* RNAPII-mut cells in each cell cycle population, and no significant differences were detected. (B) Top, diagram of the Violet Trace experimental design. Bottom, histogram overlays showing Violet Trace dilution dynamics across cell divisions. For normalization, the number of cells corresponding to the centre of each peak was considered as 100% of the cell population for that time point, thus resulting in every peak displaying the same height. (C) Left, representative images of cells in different S phase substages according to their pattern of EdU incorporation (10 μM EdU pulse during 20 min). Scale bar 5 μm. Right, histogram plots displaying the percentage of early, mid and late S substages. The number of cells in the different S phase substages was normalized to the total number of EdU+ cells. Data are from 3 independent experiments. Mean and SD are shown. Statistical significance was analysed by a Student t-test comparing WT *versus* RNAPII-mut cells between S phase substages distributions, and no significant differences were detected. (D) Top, representative Western blot membranes showing the levels of the indicated proteins in whole cell extracts from WT and RNAPII-mut cells. Aphidicolin (Aph) treatments were used as positive controls for proper DDR activation. Bottom, histogram plots showing the quantification from 3 independent experiments. Average and SD are shown. For every condition, data has been normalized to WT untreated (Unt). Statistical significance was assessed by a Student t-test to compare between WT and RNAPII-mut untreated conditions, and a 1-way Anova to compare between Unt and Aph treatments in WT and RNAPII-mut cells. Significant p-values (<0.05) are shown on top. (E) Left, representative images of γH2AX immunostaining and EdU incorporation patterns. Scale bar 5 μm. Right, γH2AX signal in early, mid and late S-phase cells sorted according to their EdU pattern. Data shown are from 1 representative experiment. Medians are indicated. Kruskal-Wallis test was used to assess statistical significance between WT and RNAPII-mut cells in each cell cycle phase and among phases. Significant p-values are indicated. (F) Top, representative images of γH2AX immunostaining in untreated cells (Unt) and upon aphidicolin (0.5 μM Aph) or 8 Gy γ-rays exposure (Irr). Scale bar 5 μm. Bottom, dot plots showing γH2AX nuclear intensity quantification. Data are from 3 independent experiments for untreated conditions, 2 independent experiments for Aph treatments and a single experiment for irradiation. n>161 cells per condition. Statistical significance was assessed by a Mann-Whitney test to compare between WT and RNAPII-mut untreated conditions, and Kruskal-Wallis followed by Dunn’s test was used to statistically compare between untreated and each treatment in WT and RNAPII-mut cells. P-values <0.05 appear on top. p<0.0001 refers to the comparisons between untreated and each treatment in WT and RNAPII-mut cells.

**Figure S2 (Related to Figures 1 and 2). S-phase progression and abundance of active RNAPII complexes on chromatin in WT and RNAPII-mut cells.** (A) Representative flow cytometry dot plot showing S-phase progression upon synchronization by double thymidine (Thy) block and release in WT and RNAPII-mut cells. Cells were pulse-labelled with 10 μM EdU for 30 min before harvesting. Asynchronous growing cells (Asyn) are shown as controls. (B) Bar graphs depicting the frequency of cells in each cell cycle phase along time. For quantification, S-phase has been divided into early (1st half) and late (2nd half) sub-stages. The top line of each bar indicates the percentage of cells at each phase as the average of 3 independent experiments. SD is shown. (C) Left, representative images of stretched DNA fibers in early (1 h), mid (3.5 h) and late (6 h) S phase. Time refers to the number of hours after release from the second thymidine (Thy) block in experiments analogous to the one shown in A. Scale bar 10 μm. Right, dot plots depicting the fork rate (FR) measured by DNA fiber analysis at the indicated time points. Data shown are from a single representative experiment (n>86 structures/condition). Kruskal-Wallis followed by Dunn’s multiple comparisons test was used to compare between groups. P-values <0.05 are shown on top. (D) Left, mid sections of 3D-SIM images representative of early and mid S phase WT cells, classified according to their EdU pattern, together with the RNAPII S2P foci. Scale bar 4 μm. Right, dot plot showing the number of RNAPII S2P foci per nucleus in WT and RNAPII-mut cells. Medians for each experiment (n=3) and total median are shown. Student t-test was used to assess statistical significance between the indicated pairs, and p-values <0.05 appear on top. Number of nuclei: WT early n=82, RNAPII-mut early n=79, WT mid n=44, RNAPII-mut mid n=41. (E) Representative Western blot membranes showing the levels of the indicated proteins in the chromatin fraction from WT and RNAPII-mut cells in synchronization experiments analogous to the one shown in A.

**Figure S3 (related to Figure 3). Differential expression analysis of coding genes between RNAPII-mut *vs* WT cells.** (A) Volcano plot showing the log_2_(fold-change) and -log_10_(p-value) of each transcript in the RNAPII-mut *vs* WT cells comparison (n=13254). Significant down-(n=393) and up-regulated (n=306) genes are indicated in purple and orange, respectively. (B) Gene size distributions of deregulated genes (DEGs) between RNAPII-mut and WT cells. (C) Normalized expression level distributions of RNAPII-mut *vs* WT DEGs. (D) Main GO-terms assigned for down-(left) and up-(right) differentially expressed genes between RNAPII-mut versus WT cells identified by clusterProfiler R package in the ontology category of biological processes. (E) Expression analysis of naive pluripotency markers over the dedifferentiation time-course. Heatmaps show the log_2_(fold-change) values between the indicated time points upon 2i exposure in days (d1, d2, and d7) versus the starting time point (d0) in WT and RNAPII-mut cells. (F) Expression levels in RPKMs of RRM1 and RRM2 at the indicated time points of the de-differentiation time-course.

**Figure S4 (related to Figure 4). Alternative splicing analyses.** (A) Mathematical formula for PSI calculation of events annotated in mm10 VASTDB library. Only events with PSI values between 10% and 90% were classified as alternative. (B-D) Differential PSI events between RNAPII-mut and WT from this work dataset and Maslon *et al* (2019) dataset (B), naïve-pluripotency and primed-pluripotency stages from our dataset (C), and additional datasets (D). (E) Correlation of changes in gene expression and AS (exons and intron retention events) in naïve-pluripotency as compared to primed-pluripotency, in WT (top) and RNAPII-mut (bottom). The y□axis represents the log_2_(fold-change) in expression, while the x□axis indicates the ΔPSI. Blue dots indicate significant genes that change in expression levels (DEGs), yellow dots show the presence of significant differentially spliced events in the represented genes, red dots represent both differentially expressed and spliced genes, and grey dots show genes that do not statistically change in their expression nor splicing pattern between conditions. The relation between these two variables is represented by the regression line and its corresponding Pearson correlation coefficient.

**Figure S5 (related to Figure 5)**. **Analyses of temporally differential DEGs.** (A) Hierarchical clustering based on the normalized expression levels (Z-scores) of genes that are differentially regulated (DEGs) in at least one naïve-pluripotency versus primed-pluripotency comparison in WT cells (d1-d0, d2-d0 or d7-d0) (n=1878 genes). Four main clusters of genes were obtained, whose expression trajectories from d0 to d7 are shown in Fig 5. Cluster 1 (gradual decrease) = 517; cluster 2 (rapid decrease) = 460; cluster 3 (rapid increase) = 267; cluster 4 (gradual increase) = 634. (B-C) Main biological functions identified by *Panther* software associated with the set of genes included in each of the four WT clusters defined in A for WT cells (C) or RNAPII-mut cells (D). Bars represent the -log_10_(p-value) of the enrichment.

## MATERIALS AND METHODS

### Cell culture and treatments

WT and RNAPII-mut E14 Mouse Embryonic Stem Cells (kindly provided by Dr. Javier Cáceres), were grown in Dulbecco’s Modified Eagle’s Medium (DMEM) supplemented with 10% FBS (Gibco), 1x non-essential aminoacids, 1 mM sodium piruvate, 2 mM L-glutamine, 50 µM β-mercaptoethanol, 100 U/mL penicillin-100 μg/mL streptomycin (Invitrogen) and 10^3^ U/mL ESGRO mLIF medium supplement (Millipore) at 37°C and 5% CO_2_ on 0.1% gelatine-coated plates. Naive pluripotency state was obtained by adding 1 µM PD0325901 (MEK inhibitor; StemMACS Miltenyi Biotec, #130-103-923) and 3 µM CHIR99021 (GSK3 inhibitor; StemMACS Miltenyi Biotec, #130-103-926) to the same medium and growing the cells for 7 days, changing the media every day and splitting the cells every other day. Cells were routinely tested for mycoplasma contamination.

Aphidicolin (Aph, Sigma Aldrich A0781) treatments were performed at two concentrations (0.1 and 0.5 µM) and incubation times (2.5 and 16 h), as indicated.

Cells were synchronized at the G1/S border by double thymidine (Thy; Sigma Aldrich T1895, SLCG7664) blocking as follows: 2 mM Thy was added to the culture media for 16 hours, cells were then washed twice with pre-warmed PBS and released for 8 h into Thy-free medium followed by another 16h incubation with 2 mM Thy.

### Cell cycle and cell proliferation analysis

For cell cycle analysis, cells were pulse-labelled with 10 µM EdU for 20 min, harvested by trypsinization, fixed with 4% PFA in PBS for 15 min at RT and permeabilized with 0.25% Triton X-100 for 20 min at RT. EdU incorporation was revealed by Click-it reaction for 30 min. DNA was stained with DAPI in the presence of 0.25 mg/mL RNase A/T1 (Thermo Scientific, EN0551). Samples were processed in a FACSCanto II cytometer (Becton Dickinson) with FACSDiva v6.1.3 analysis software and data was analyzed with FlowJo v10 software (Tree Star Inc).

Cell proliferation was analysed by CellTrace^TM^ Violet (Invitrogen #C34571) according to manufacturer’s instructions. Briefly, harvested cells were labelled in suspension with 5 µM CellTrace^TM^ Violet dye for 20 min at 37°C. After 2x wash with culture medium with FBS, an equal number of cells were plated to be collected every 12 h. Samples were fixed and processed as cell cycle analysis samples.

### DNA fiber Stretching

Exponentially growing cells were sequentially pulse-labelled with 50 µM CldU (Sigma; #C6891) and 250 µM IdU (Sigma; #I5125) for 20 min each, trypsinized and collected in cold PBS at a concentration of 2.5x10^5^ cells/mL. 2 µL of cell suspension were lysed by the addition of 10 µL of pre-warmed (30°C) spreading buffer (200 mM Tris pH 7.4, 50 mM EDTA, 0.5% SDS) on top of a glass slide. After 6 min incubation in a humidity chamber at RT, slides were tilted on a 30° angle to stretch the DNA. Fibers were air dried and fixed with 3:1 ice-cold methanol:acetic acid solution for 2 min, denaturalized with 2.5M HCl solution for 30 min at RT and treated with blocking solution (1% BSA, 0.1% Triton X-100) for 1h. For immunodetection of labeled tracks, samples were incubated with 1:100 anti-CldU (Abcam, Ab6326) and 1:100 anti-IdU (BD Biosciences, 347580) antibodies at 4°C overnight, followed by 1h incubation at RT with 1:300 secondary antibodies (anti-rat IgG Alexa Fluor 647; ThermoFisher, 347580 and anti-mouse IgG1 Alexa Fluor 488; ThermoFisher A21121). A third incubation was performed with 1:200 anti-ssDNA (Millipore, MAB3034) and 1:300 anti-mouse IgG2a Alexa Fluor 555 (ThermoFisher, A21137) antibodies, simultaneously for 30 min at RT. Following air-drying, slides were mounted with ProLong Glass AntiFade (ThermoFisher; #P36984).

Images were acquired using a 63X/1.4 oil plan-apochromatic objective in an Axiovert200 Fluorescence Resonance Energy Transfer Zeiss microscope and Metamorph 7.10.4.407 software. Analysis of track lengths and origins of replication was performed using FIJI/ImageJ version 2.14.0. Calculation of fork rate was done based on the conversion factor 1 µm=2.59 kb (Jackson and Pombo, 1998). Statistical analysis was carried out in Prism v8 (GraphPad Software) using the non-parametric Mann-Whitney rank sum or Kruskal-Wallis tests.

### Protein extraction and western blotting

Whole protein extracts were obtained with a urea-based solution (8 M urea, 50 mM Tris-HCl pH 8, 1% CHAPS). Cell pellets were resuspended in 2 to 3 volumes of extraction buffer and incubated for 30 min at 4°C in agitation. After centrifugation at maximum speed for 5 min at 4 °C, proteins in the soluble fraction were collected and their concentration assessed by Bradford assay. Biochemical fractionation was performed as described in Méndez and Stillman (2000).

SDS-PAGE, protein transfer to a nitrocellulose membrane (Amersham^TM^ Protran^TM^ 0.45 µm) and immunoblotting were performed following standard protocols. Visual acquisition was done in an Amersham 680-RGB and band intensities were measured using FIJI/ImageJ software. Antibodies used are listed in Table S1.

### Immunofluorescence and nascent DNA labelling

Cells grown on gelatine-coated glass coverslips were pulse-labelled with 10 µM EdU (Merck, 900584) during 20 min for conventional and QIBC microscopy, and 5 min for 3D-SIM. For the 3D-SIM experiments, after EdU pulse, cells were washed with pre-warmed PBS and incubated with pre-warmed medium for 5, 20, 40 and 70 minutes of chase before fixation. Cells were fixed with 4% PFA (Electron Microscopy Sciences) in PBS for 15 mins and permeabilized with 0.25% Triton X-100 for 20 min at RT. Blocking was carried out with 3% BSA (Sigma) overnight at 4°C for conventional confocal imaging and with MaxBlock (Active Motif) for 30 min at RT for 3D-SIM and QIBC microscopy, before Click-it reaction and/or primary antibodies incubation.

Click-it reaction cocktail (5 mM CuSO4, 10 % Tris-HCl pH 8, 50 mM sodium ascorbate) for EdU detection was done using 0.5% Azide Fluor 488 (Jena Bioscience, CLK1275) for conventional microscopy and 0.25% Azide Fluor 549 (Thermo Fisher, A10270) for 3D-SIM. Primary antibodies were incubated overnight at 4 °C and secondary antibodies were incubated for 1 h at room temperature (Table S1). For 3D-SIM imaging, samples were post-fixed with 4 % PFA for 10 min. Coverslips were counterstaining with DAPI (1:1500, Millipore 5 mg/mL) and mounted with Prolong Glass (ThermoFisher, for conventional microscopy), Mowiol (Sigma Aldrich, 81381-for QIBC) or SlowFade (TM Diamond Antifade Mountant Invitrogen for 3D-SIM).

For conventional confocal microscopy image acquisition, a A1R+ confocal microscope (Nikon) with 60x oil objective was used. Nuclei segmentation was automatically performed by CellPose Advanced ImageJ plugin based on DAPI staining, and nuclear intensity/foci number of the different targets was analysed using FIJI/ImageJ.

### 3D Structured-Illumination Microscopy (3D-SIM)

For Super-Resolution, imaging was performed on a DeltaVision OMX SR microscope, using a 60X oil immersion objective. Computational image reconstructions were done using channel specific OTFs (Optical Transfer Functions) with softWoRx 7.2.0. Prior to the analysis, processing and quality control of the reconstructed images was carried out using SIMCheck in FIJI. Processing comprised 16-bit conversion and thresholding, and quality control consisted of a Modulation Contrast Map-based filtering, as well as a manual registration of the images. Selected images were then channel-aligned using Chromagnon (v0.92). Analysis was done using a FIJI custom-written macro to calculate the percentage of overlapping between DNA being synthesized (EdU) and elongating transcription (RNAPII S2P). Briefly, for each RNAPII S2P foci, its brightest point (centroid) was detected and considered as an “overlapping” foci if co-localizing with an EdU pixel. In parallel, the total number of RNAPII S2P foci per nucleus was quantified, as well as nuclei volume and DNA content based on DAPI staining. Cells were manually classified in early or mid S phase according to their EdU pattern.

### Quantitative Image-Based Cytometry (QIBC)

QIBC experiments for detection of γH2AX along the cell cycle were performed on an Olympus ScanR inverted widefield microscope system equipped with IX83 inverted motorized frame with Z-drift control, and Semrock DAPI/FITC/Cy3/Cy5 Quad LED filter set. Light sources were Lumencor SPECTRA X Light Engine Independent LEDs and an Olympus UPLXAPO 20x Air objective was used.

Images were captured using SCANR Acquisition software (Version 3.2.0) and analysed, including nuclei segmentation, using the Olympus ScanR Image Analysis Software (Version 3.2.0). Cell cycle phase was addressed based on EdU and DAPI mean intensities in each cell. Data from the analysis was visualized using TIBCO Spotfire software (Version 12.5.0).

### Total RNA extraction and sequencing

Cells were resuspended in TRIzol^TM^ (ThermoFisher) and total RNA was subsequently isolated by isopropanol precipitation. RNA samples were treated with DNAse using Kit TURBO DNA-free^TM^ (Invitrogen) following manufacturer’s instructions.

For total RNA-seq, libraries were prepared using the DNBSEQ Eukaryotic Strand-specific mRNA kit (Illumina) at BGI Tech Solutions and sequenced on an Illumina DNBSEQ-G400 Sequencing Platform to produce 150-nucleotide long paired-end reads at 30M coverage with a phred quality score Phred+33.

### RNA-seq transcriptomic analyses of protein coding genes

Sequenced reads were evaluated for quality using FastQC (Andrews, 2010) and aligned to the mm10 reference genome using *STAR* (Dobin et al., 2013) with standard parameters. BigWig files loaded in the IGV browser (Robinson et al., 2011) were generated with the *bamCoverage* function from *deepTools* (Ramírez et al., 2016) using parameters *-normalizeUsing* CPM and *-bs* 1. Expression levels in the protein coding transcriptome were computed with *featureCounts* (Liao et al., 2014) using the RefSeq annotation (O’Leary et al., 2016) for the longest transcript variant, and normalized by the total read number in each experiment and the gene size.

Differential gene expression analysis between conditions was performed with *DESeq2* (Love et al., 2014). Genes showing adjusted p-value < 0.05 and |log_2_(fold-change)| > 0.5 were considered as significantly differentially expressed (DEGs). GO term enrichment analyses of RNAPII-mut *vs* WT DEGs at d0 were computed using the *enrichGO* function from R *clusterProfiler* package (Yu et al., 2012) with default parameters and the “*BP*” (biological processes) ontology category. GO term analysis of UMAP genes clustered according to their trajectories along de-differentiation were performed using *Panther* v19.0 (Mi et al., 2019), and the top ten listed genes, sorted by the smallest FDR value, were selected for representation.

Rank–rank hypergeometric overlap (RRHO) test was performed using the ranked list of log_2_(fold-change) values in the most naïve (d7) *vs* most primed (d0) stages, in WT and RNAPII-mut cells. Heatmap colours represent the -log_10_(p-value) after Benjamini-Yekutieli correction of the hypergeometric overlap analysis (Plaisier et al., 2010).

Hierarchical clustering of DEGs in at least one naïve versus primed comparison in WT (d1-d0, d2-d0 or d7-d0) (n=1878 genes) was performed based on their normalized expression levels (Z-scores). Uniform Manifold Approximation and Projection (UMAP) algorithm of these genes was used to visualize their similarities in terms of expression changes along the primed-to-naive transition using the *umap* package from R (Konopka, 2025). For that, we used their normalized expression levels (Z-scores) in all time-points (d0, d1, d2, and d7). Each UMAP dot represented a gene, which was coloured depending on its cluster classification in the hierarchical clustering analyses or its log_2_(fold-change) value in the RNAPII-mut vs. WT differential expression analyses.

Heatmaps and histone metagene profiles were performed using, respectively, *pheatmap* (Kolde, 2019) and *metagene2* (Fournier et al., 2024) packages from R. Both violin and boxplots were generated using *ggplot2* package from R (Wickham, 2016).

### Alternative splicing analysis

Alternative splicing (AS) analysis from total RNA-seq samples was performed using *vast-tools* v2.5.1 (Tapial et al., 2017) with mm10 mouse VASTDB library (vastdb.mm2.23.06.20.tar.gz). The analyzed event types included “exon,” “intron,” “alternative 3’ splice site,” and “alternative 5’ splice site”. We classified “ANN,” “S,” “C3,” “C2,” “C1,” and “MIC” as exon events. To enhance resolution, we applied the *merged* function to both replicates in each condition and defined AS events as those with PSI values between 10% and 90%. Pairwise differential AS analysis was performed using the *compare* function with parameters *--min_dPSI* 15 and *--min_range* 5.

## Data availability

Total RNA-seq data generated and analyzed in this work have been deposited in the NCBI Gene Expression Omnibus database (Barrett et al., 2013) under the accession number GSE293593.

The following GEO and ArrayExpress datasets were analyzed in this study: RNA-seq from WT mESCs in serum/LIF and serum/LIF+2i (GSE112208), RNA-seq from mouse embryos at E3.5 and E4.5 stages (E-MTAB-2958), single-cell RNA-seq from mid and late blastocyst stages (GSE45719), and RNA-seq from WT and RNAPII-mut (slow/slow) mESCs (GSE127741).

Marker gene sets for cell cycle stages were obtained from https://github.com/hbc/tinyatlas/blob/master/cell_cycle/Mus_musculus.csv, which is based on gene lists described by Tirosh et al. (2016).

## Author contribution

## Supporting information

Figure S1_Martin-Virgala et al

Figure S2_Martin-Virgala et al

Figure S3_Martin-Virgala et al

Figure S4_Martin-Virgala et al

Figure S5_Martin-Virgala et al

