## Supplementary figures and images for "Slow RNAPII elongation enhances naïve-pluripotency rewiring while preserving replication fork speed"

### Figure S1_Martin-Virgala et al

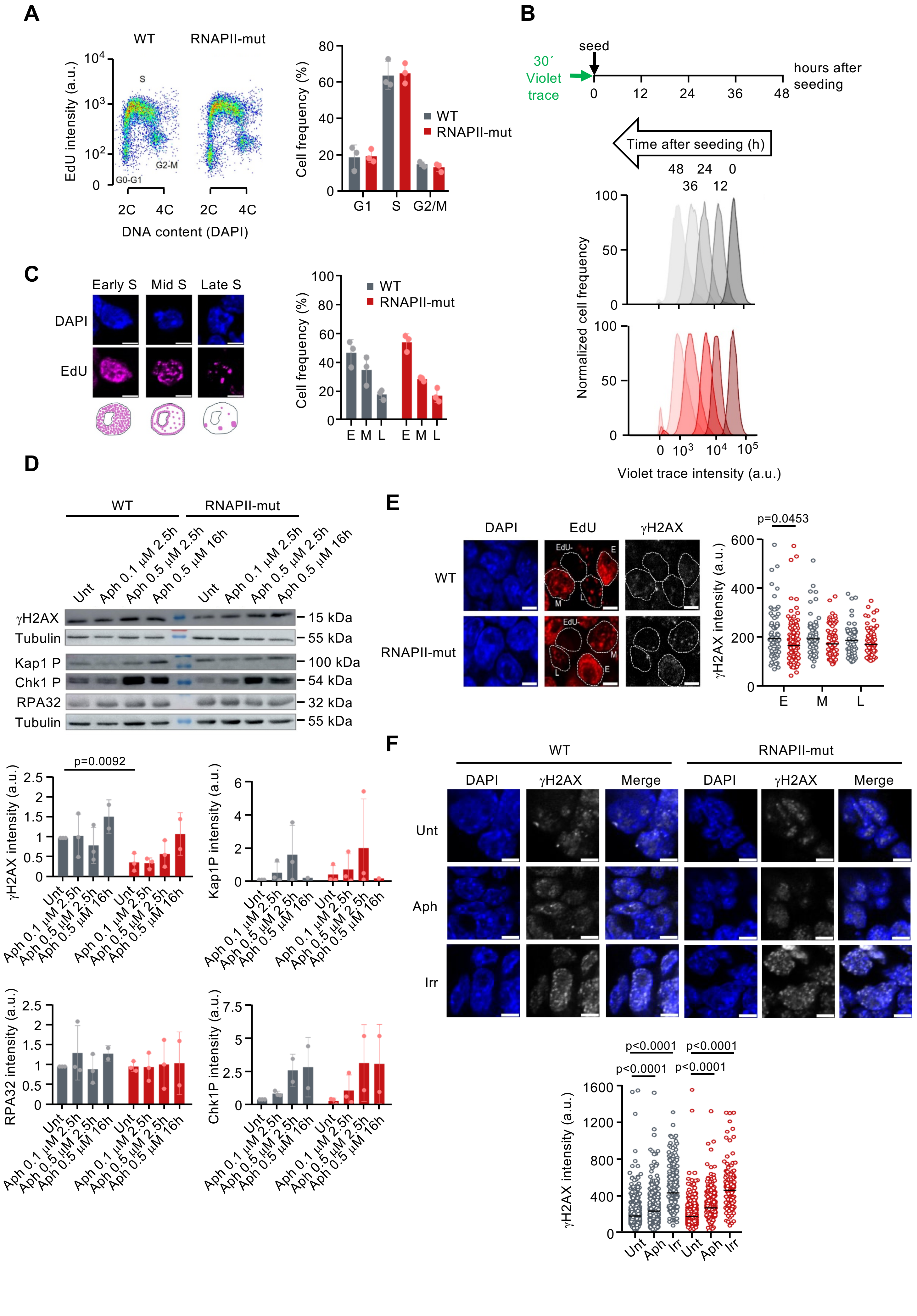

### Figure S2_Martin-Virgala et al

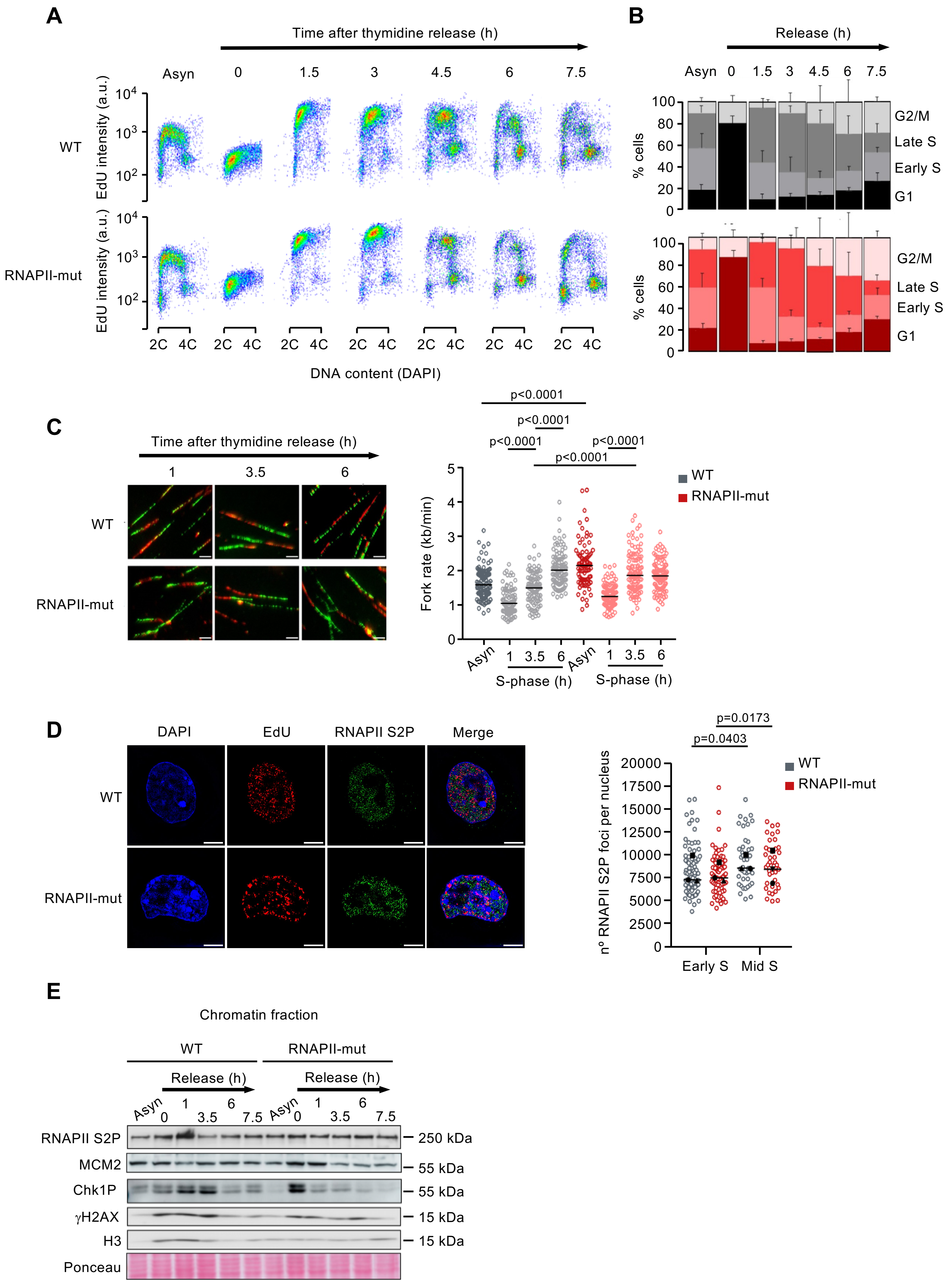

### Figure S3_Martin-Virgala et al

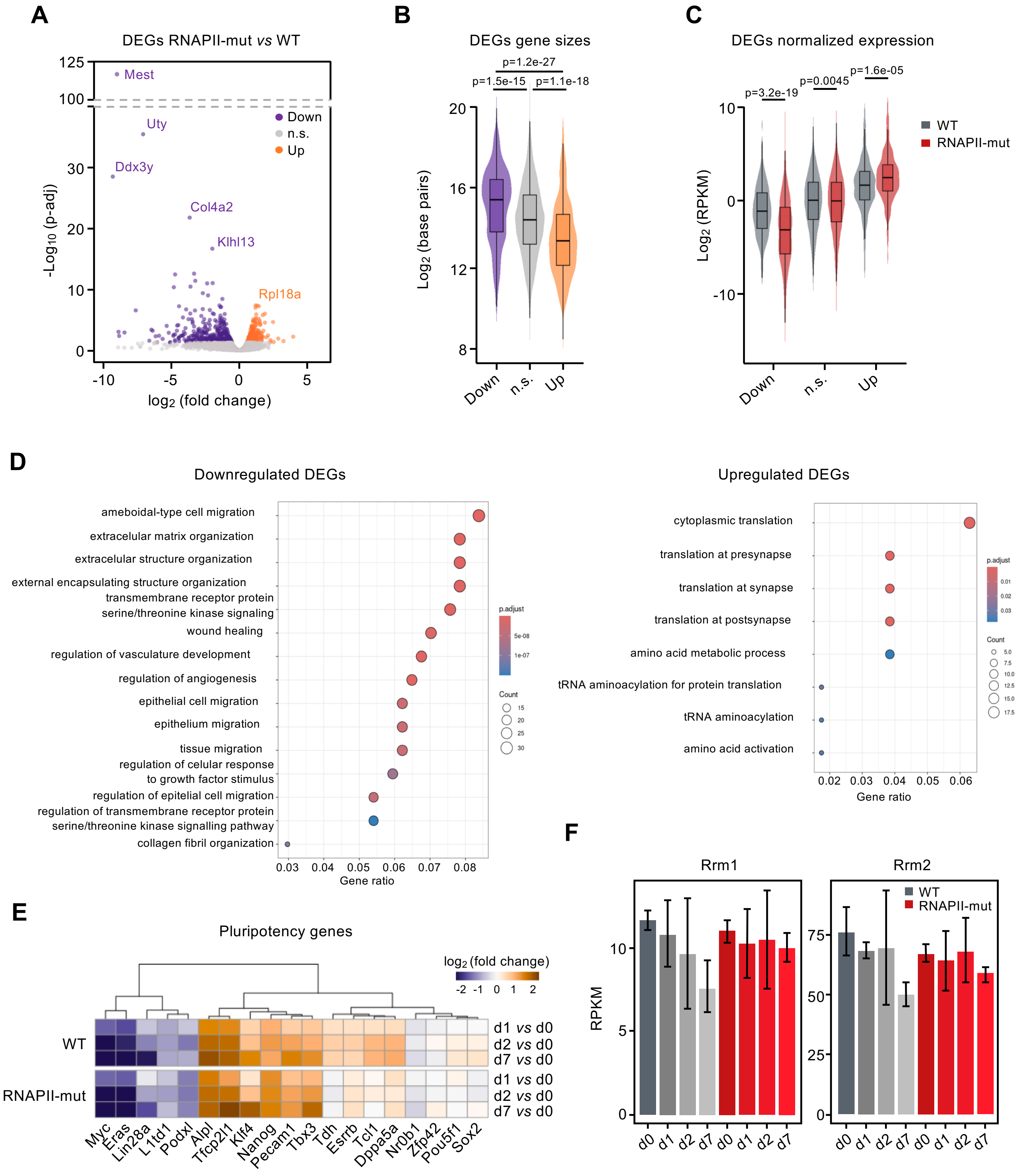

### Figure S5_Martin-Virgala et al

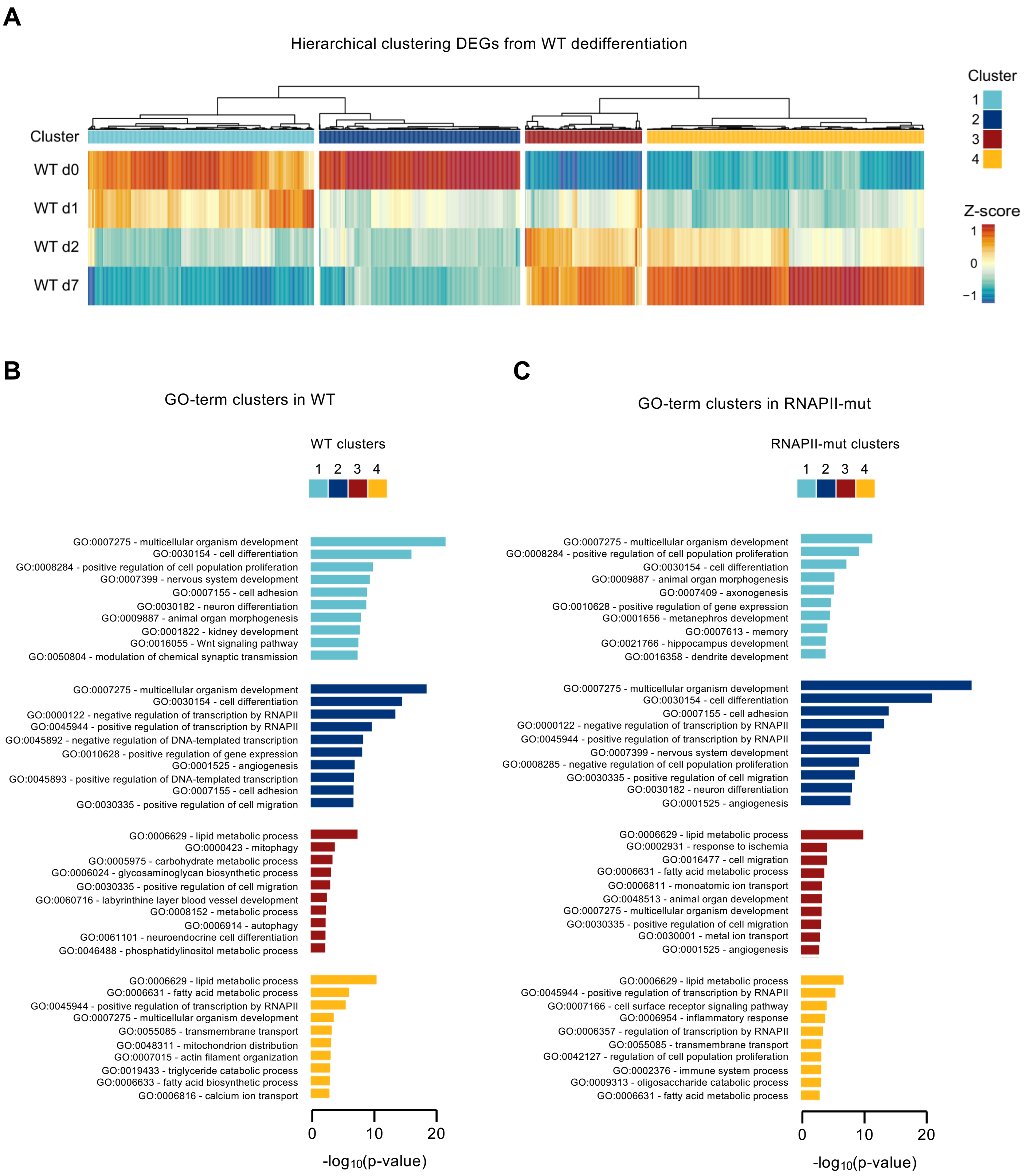
