## Supplementary material for "Slow RNAPII elongation enhances naïve-pluripotency rewiring while preserving replication fork speed": Figure S4_Martin-Virgala et al

**A**

Total AS in each condition

$$PSI = \frac{\text{Number of reads supporting the inclusion of an event}}{\text{Total number of reads (including both inclusion and exclusion)}}$$

- Cryptic events: PSI < 10%
- Alternative spliced events: PSI 10-90%
- Constitutive events: PSI > 90%

**B**

ΔPSI between RNAPII-mut and WT
(own data and other datasets)

| RNAPII-mut vs. WT | own data | Maslon *et al.* data |
|---|---|---|
| Exons in WT | 357 | 316 |
| Exons in RNAPII-mut | 222 | 210 |
| Introns in WT | 109 | 176 |
| Introns in RNAPII-mut | 318 | 243 |
| Alternative 3’ ss | 173 | 167 |
| Alternative 5’ ss | 158 | 198 |
| Total | 1337 | 1310 |

**C**

ΔPSI events between naïve-pluripotency (day 7)
and primed-pluripotency (day 0) stages
(own dataset)

| naive *vs* primed | WT | RNAPII-mut |
|---|---|---|
| Exons in day 0 | 198 | 151 |
| Exons in day 7 | 391 | 389 |
| Introns in day 0 | 169 | 267 |
| Introns in day 7 | 235 | 105 |
| Alternative 3’ ss | 169 | 154 |
| Alternative 5’ ss | 187 | 137 |
| Total | 1349 | 1203 |

**E**

Correlation between expression and splicing

WT (day 7 *vs* day 0)

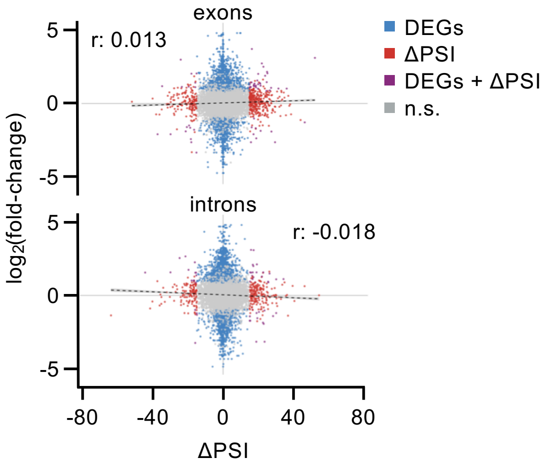


RNAPII-mut (day 7 *vs* day 0)

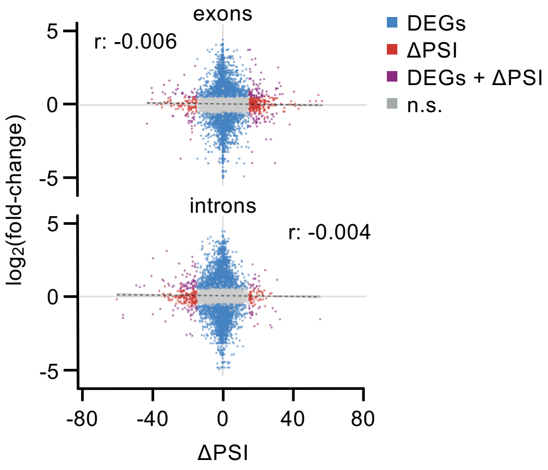


**D**

ΔPSI events between naïve-pluripotency and primed-pluripotency stages
(other datasets)

| naive *vs* primed | mESCs (Lynch *et al.*) | mid *vs* late-blastocyst (Deng *et al.*) | E3.5 *vs* E4.5 embryo (Boroviak *et al.*) |
|---|---|---|---|
| Exons in primed pluripotency | 161 | 762 | 874 |
| Exons in naive pluripotency | 232 | 1212 | 932 |
| Introns in primed pluripotency | 76 | 1952 | 668 |
| Introns in naive pluripotency | 82 | 2515 | 909 |
| Alternative 3’ ss | 138 | 733 | 752 |
| Alternative 5’ ss | 110 | 595 | 504 |
| Total | 799 | 7769 | 4639 |

Figure S4
